# Using bootstrap procedures for testing the modular partition inferred via leading eigenvector community detection algorithm

**DOI:** 10.1101/2021.10.04.462969

**Authors:** Oksana Vertsimakha, Igor Dzeverin

**Affiliations:** Taras Shevchenko National University of Kyiv, Kyiv, Ukraine; Schmalhausen Institute of Zoology, Kyiv, Ukraine

**Keywords:** Modularity, Community detection, Leading eigenvector algorithm, Networks

## Abstract

Modularity and modular structures can be recognized at various levels of biological organization and in various domains of studies. Recently, algorithms based on network analysis came into focus. And while such a framework is a powerful tool in studying modular structure, those methods usually pose a problem of assessing statistical support for the obtained modular structures. One of the widely applied methods is the leading eigenvector, or Newman’s spectral community detection algorithm. We conduct a brief overview of the method, including a comparison with some other community detection algorithms and explore a possible fine-tuning procedure. Finally, we propose an adapted bootstrap-based procedure based on Shimodaira’s multiscale bootstrap algorithm to derive approximately unbiased p-values for the module partitions of observations datasets. The proposed procedure also gives a lot of freedom to the researcher in constructing the network construction from the raw numeric data, and can be applied to various types of data and used in diverse problems concerning modular structure. We provide an R language code for all the calculations and the visualization of the obtained results for the researchers interested in using the procedure.

## 1 Introduction

Modularity and integration are of great interest for evolutionary biologists. Modular structure can be recognized at multiple levels of biological organization, and it is assumed to reflect evolutionary or developmental processes. Various methods based on the analysis of patterns found in trait correlation or covariance matrices are now available (Klingenberg, 2008; Goswami & Polly, 2008), as well as various measures of the level of modularity itself (Eble, 2005; Adams et al., 2004; Mitteroecker, 2009). At the same time, modularity and community detection are fundamental problems of the modern networks theory (Freeman, 2004). In recent years, in many fields of biology networks and network analysis methods have found numerous implementations (Pilosof et al., 2017; Thébault & Fontaine, 2010; Mahmoud et al., 2013). For example, in spatial ecology studies of species as parts of communities or networks came into focus (Landi et al., 2018; Staniczenko et al., 2017; Ovaskainen et al., 2017). Various types of networks, such as ecological, phylogenetic, neurological, metabolic or molecular interaction networks arise in studies on different levels (Barabasi et al., 2011; da Fontura Costa et al., 2008; Newman, 2003). Network theory methods have given rise to some state-of-the-art research approaches such as anatomical networks analysis (AnNA) (Esteve-Altava et al., 2015; Diogo et al., 2021; Ostachuk, 2019), and have also become valuable addition to some of the traditional methods, including morphological studies of integration and modularity. In those studies, one of the crucial questions concerns the statistical support of the obtained structure (Shimodaira, 2004). We take a closer look at one of the network community detection methods known as Newman’s spectral, or the leading eigenvector, algorithm: even though this algorithm has already been applied in various studies (Brederoo et al., 2021; Xu et al., 2021; Labatut & Balasque, 2012; Xia et al., 2013; Hagmann et al., 2008) to the best of the authors knowledge, no comprehensive analysis has been conducted yet. Therefore, we include a brief overview of the algorithm and its comparison to some other widely used community detection algorithms on synthetic benchmarks as well as a real-world networks collection.

While various methods have been developed in recent years, there exist definite theoretical limits of the community detection. No algorithm can be optimal on all kind of inputs, and so the methods efficacy would differ on different data; on the other hand, there is no bijection between network structure and ground truth communities, and therefore no algorithm can always recover the correct ground truth on every network (**??**). Therefore, the problem of estimating statistical significance of the obtained partitions becomes highly important.

We propose a procedure that allows to model a network based on the correlation matrix of the traits. Approximate p-values are computed for all the subdivisions suggested by the algorithm with the Shimadaira’s multiscale bootstrap method (Shimodaira, 2004). With our code the results can also be visualised and summarized to help researchers to make an informed decision on the final partition onto modules. We propose various options for the model network construction including the implementation of the fine-tuning procedure for the leading eigenvector algorithm proposed in the original article.

## 2 Materials and Methods

We focus on the leading eigenvector algorithm introduced in (Newman, 2006b,a). The algorithm is based on the modularity maximization. If we consider an undirected graph *G* = (*V, E*) with vertices 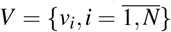 and respective degrees 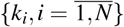 and with adjacency matrix 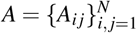, then modularity is defined as

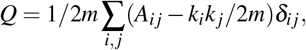

where 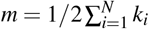 is the total sum of the degrees *δ*_*i j*_ = 1 if *v*_*i*_ and *v* _*j*_ belong to the same module, and 0 otherwise.

A split into two modules can be given as a vector 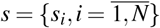 where *s*_*i*_ *ϵ*{1, −1}, and thus, the definition can be rewritten as

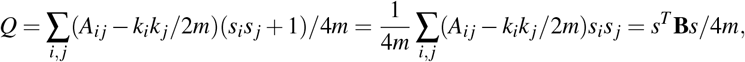

for *B*_*i j*_ = *A*_*i j*_ − *k*_*i*_*k* _*j*_/2*m*.

Now, if we denote the eigenvalues of the matrix **B** (sorted in descending order), with the corresponding eigenvectors 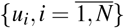 and 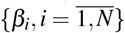 respectively, modularity of the split *s* equals:

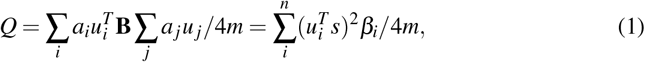

where *a* denotes the coordinates of the *s* given in the eigenbasis. The eigenvectors of **B** are non-negative (Newman, 2006b), therefore, we consider vector *s* where *s* _*j*_ = 1 if the corresponding coordinate of the *u*_1_ > 0 and *s* _*j*_ = 0 otherwise, 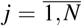 To make further splits, we calculate the increment of the modularity by substituting **B** in (Eq. 1) with

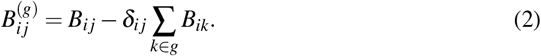

The component of the network is further subdivided as long as new partition results in increase in modularity. Each split has *O*((*N* + |*E*|)*N*) complexity (it becomes *O*(*N*^2^)on a sparse network). In practice, the algorithm rarely splits all the nodes, and the average running time for the whole algorithm can be estimated as *O*(*N*^2^*log*(*N*)) (Newman, 2006b; Mahmoud et al., 2013). Community detection algorithms selected for comparison are two modularity-based algorithms with complexity of *O*(|*E*|), Louvain (Blondel et al., 2008) and Fast Greedy (Rosvall & Bergstrom, 2007; Z. Yang et al., 2016), Infomap, an information flow based algorithm with complexity *O*(|*E*|) (Rosvall et al., 2010; Raghavan et al., 2007; Z. Yang et al., 2016), and Walktrap algorithm (complexity of *O*(|*E*|*N*^2^) or *O*(*N*^2^*log*(*N*) for sparse networks) based on random walks (Pons & Latapy, 2006). For all of the algorithms including the basic leading eigenvector algorithms, we used implementations included in the R language igraph 0.9.2 package (Csardi & Nepusz, 2005). The fine-tuning procedure for the leading eigenvector is based on the Kernighan-Lin algorithm (Kernighan & Lin, 1970) and was proposed and described in the original paper (Newman, 2006b). We implemented it as an R language function, the code and the detailed description of the functions is available in the github repository (see Data Availability). To measure the similarity between different partitions, we used normalized mutual information (Knuth, 1993) as defined in (Fred & Jain, 2003). Mean NMI values were used to assess the similarities between different methods and between obtained and meta partitions to compare results on the benchmarks. As characteristics of the partitions we used modularity and significance, an alternative measure of modular structure quality proposed and discussed in (Traag et al., 2013). To examine the differences in obtained results we used paired Wilcoxon signed rank test with continuity correction. To compare the methods we used an open collection of 191 networks from the biological domain of the real-world networks collection *CommunityFitNet* (Ghasemian et al., 2020). Further, we generated sets of benchmarks of two different types, Girvan-Newman (GN) graphs (Newman & Girvan, 2004) and LFR (Lancichinetti et al., 2008) graphs. Networks were generated with *NetworkX* 2.4 library for Python (Hagberg et al., 2008)) The results were averaged over 300 generations for each set of parameters. Each GN network consists of 128 nodes divided into 4 equal modules, with average degree *k* = 16. As long as external degree *k*_*out*_ ≤ 8, the GN graphs have a well-defined community structure, with more links inside the modules than in-between. In LFR graphs degrees and sizes of the modules follow exponential distributions that can often be observed in real-world networks, the respective degrees of the distributions are denoted as *τ*_1_ and *τ*_2_. Each node has a fraction of (1− *µ*) edges connected to the nodes of the same module. For the GN graphs, we used *k*_*out*_ parameter ranging from 1 to 10. For the LFR graphs we considered networks with 150, 500 and 1000 nodes with parameters given in Table 2; parameter *µ* ranged from 0.05 to 0.5.

**Table 1.** LFR benchmark generation parameters.

| $N = \vert V\vert $ | $\tau_1$ | $\tau_2$ | $Mink$ | $Maxk$ | $MinK$ | $MaxK$ |
| --- | --- | --- | --- | --- | --- | --- |
| 150 | 1.1 | 1.1 | 5 | 30 | 5 | 40 |
| 500 | 3 | 1.1 | 5 | 50 | 10 | 70 |
| 1000 | 3 | 1.2 | 5 | 70 | 10 | 100 |

**Table 2.** Mean NMI between the methods on the set of real-world networks

| Infomap | Louvain | Walktrap | Leading eigenvector | Fine-tuning | FastGreedy |
| --- | --- | --- | --- | --- | --- |
| 0.443 | 0.505 | 0.438 | 0.456 | 0.486 | 0.499 |

To apply the leading eigenvector to the observations datasets we built weighted graphs based on the correlation matrices. We considered several possible approaches to deal with the presence of negative correlation coefficients, with two implemented in our code: bringing all the coefficients into power (i.g. square), or using *e* ^−(1 −*C*)/*sd*(*C*)^, where *C* is the correlation matrix and *sd* is the standard deviation.

To calculate the approximately unbiased p-values, we extended the application of the Shimodaira’s multiscale bootstrap estimation (Shimodaira, 2004) available for the classical hierarchical clustering in the R language *pvclust* package, to the case of not fully resolved partitions. The estimates are calculated as follows:

Let *X* be a numeric data set with*K* communities *C*_1_, …, *C*_*K*_.

1. The scaling parameters *r*_1_, …*r*_*J*_ are specified. For each one, *B*_*j*_ bootstrap samples 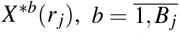 of the size *N*_*j*_ = [*Nr*_*j*_] are generated, and then the clustering algorithm is applied to each sample.
2. The fraction of times the original module was detected, *BP*(*r*_*j*_), is calculated.
3. The additional parameters *d* and *c* are estimated. For that purpose, 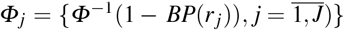 is defilned, and the weighted least squares method is applied to the vector *Φ* with vectors *r*_1_, …*r*_*J*_ and 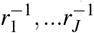 as the regressors (no intercept). The estimates for *d* and *c* are then found by minimizing residual sum of squares

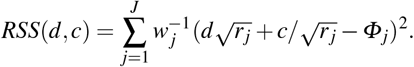 The weights used are 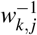 with

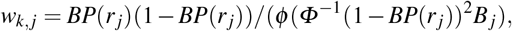

where *φ* (·) and *Φ* (·) are the quantile and density functions of the standard normal distribution respectively.
4. Approximately unbiased p-value estimations (*AU*) and classical bootstrap estimations (*BP*) are calculated as:

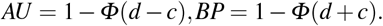

To illustrate the procedure we used a waterstrider data set that contains 8 measurements of the 6 species of the waterstrider genus *Limnoporus* Stal 1868 (Gerridae, Heteroptera, Hemiptera, Insecta). This dataset is stored in SB Morphometrics (Rohlf, 2016) and studied in (Klingenberg & Spence, 1993). We limit ourselves to the sample of the adult *L. notabilis* (Drake and Hottes 1925) at the later stages of the development (60 specimens) since the sex is unknown for earlier stages. The effects of sex and instar were excluded using a MANCOVA technique implemented in *evolQG* package (Melo et al., 2015). We also compared the output with the LModularity function output. Visual network representations and illustrations were created with *igraph* (Csardi & Nepusz, 2005), *ggplot2* (Wickham, 2016) and *corrplot* (Wei & Simko, 2017) R packages, and Gephi 0.9.2 Bastian et al. (2009) software. We used Python 3.7 and R 3.5.1.

## 3 Results

Data on the dissimilarities of various methods can be used to compare in studies methods that would typically yield contradicting results (for example, see Rahiminejad et al. (2019); Wang et al. (2015)). Analysis of the algorithm results on the real-world biological networks suggests that the leading eigenvector algorithm both with and without the fine-tuning procedure yields results most similar to those of the Louvain and FastGreedy algorithms, yet the mean NMI doesn’t exceed the value of 0.64 for any two methods. At the same time, on the simulated networks, both the basic and the fine-tuned leading eigenvectors algorithms tend to be more similar to the Walktrap algorithm and not to FastGreedy. In both cases, the most distant algorithm was Infomap. Other measures such as variance of information (Meila, 2003) or Adjusted Rand index (Hubert & Arabie, 1985) yield the same results.

We found that the fine-tuning procedure leads to significant (*p* ≤ 0.0001) improvement in the modularity maximization performance of the leading eigenvector algorithm. Figs 1–2 and Table 1 show the mean results for the real-world and the simulated networks. The results for both leading eigenvector algorithms performed better on smaller benchmarks.

**Fig. 1.**
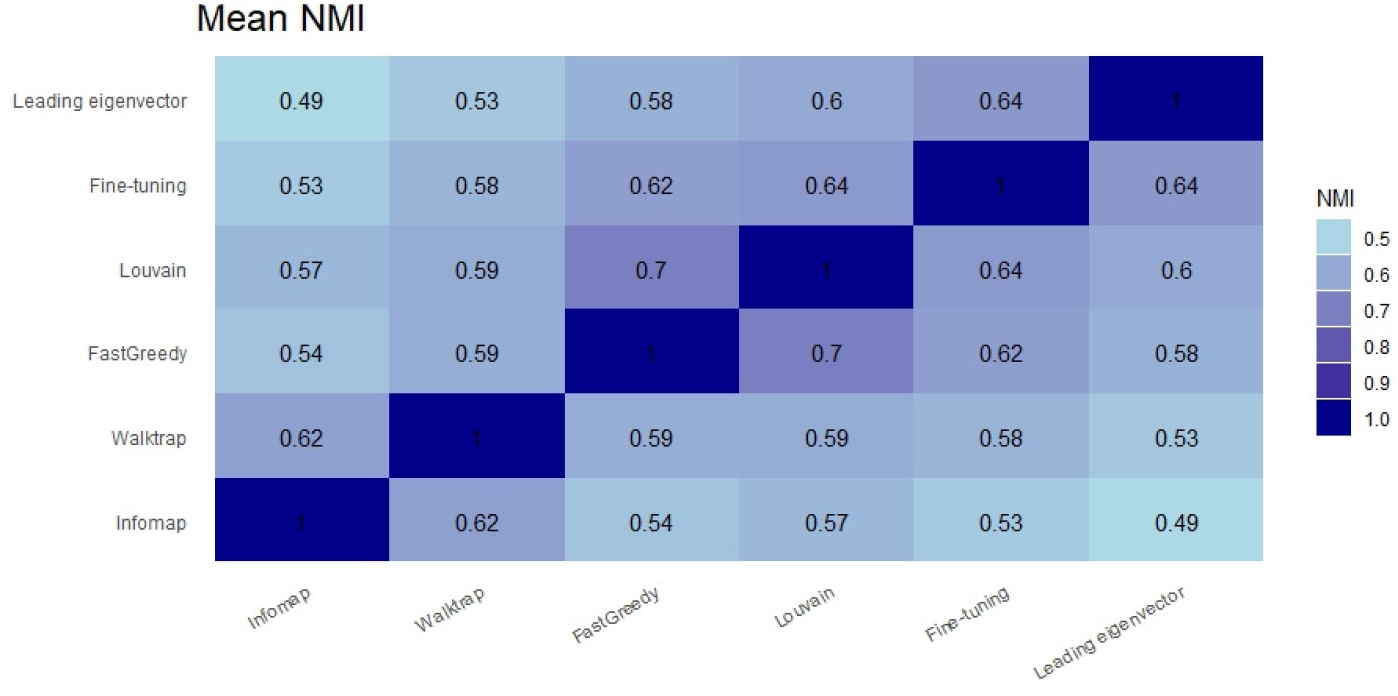
Mean NMI between the methods on the set of networks

**Fig. 2.**
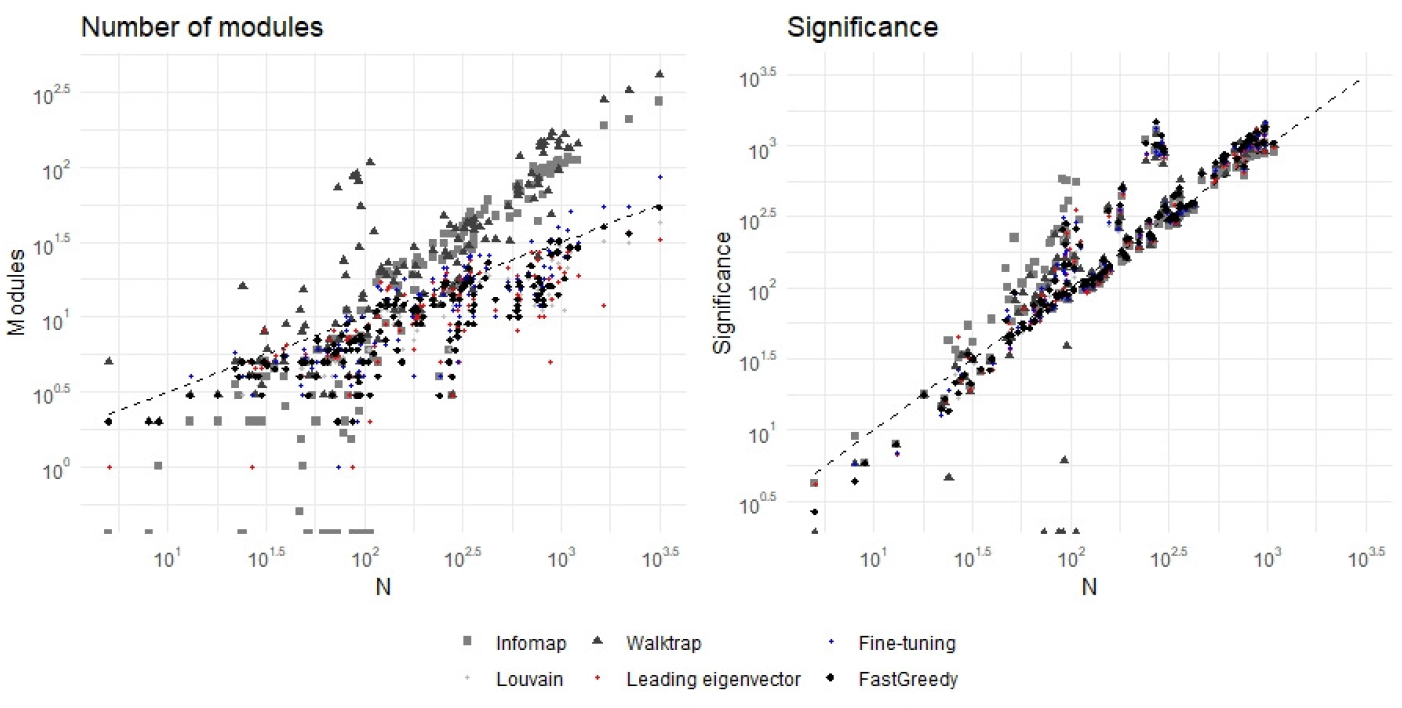
Mean number of nodes (left) and significance (right) versus number of nodes (*N*) of the *Community-FitNet* networks. Dotted lines correspond to the 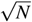 and *N* respectively.

One of the known drawbacks of the modularity maximization algorithms is the resolution limit, i.e. these methods fail to detect modules of relatively small clusters in large networks, however well defined (see Lancichinetti & Fortunato (2011); Good et al. (2010)). We compared the number of modules detected by the leading eigenvector. Several studies suggest that the maximum number of detectable communities in real-world networks is 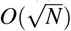 (Ghasemian et al., 2020). We examined the number of modules detected of the leading eigenvector algorithm with and without the fine-tuning procedure on the *CommunityFitNet* corpus, and found a significant increase in the number of communities (*p ≤*0.0001) after the fine-tuning procedure (Fig. 2).

Figures 3–4 show the results of the GN and LFR benchmarks, and indicate that the performance of the leading eigenvector algorithm can be improved with the fine-tuning procedure, but it still deteriorates on the large networks.

**Fig. 3.**
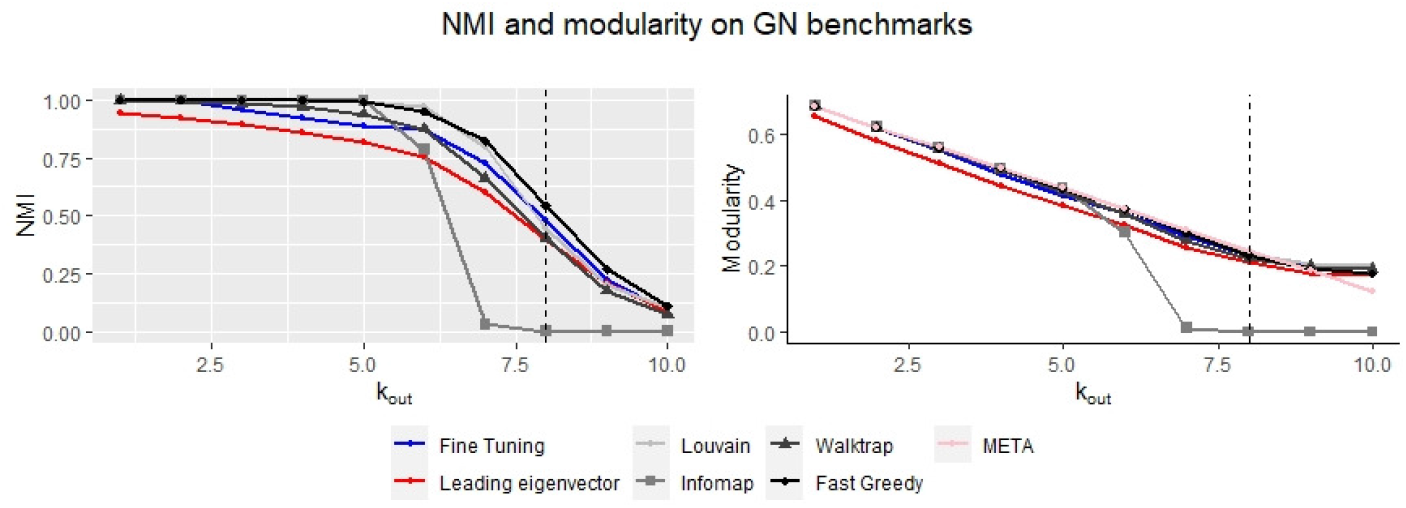
Mean modularity and NMI with planted partitions of the GN networks. Dotted line marks *k*_*out*_ = 8 point.

**Fig. 4.**
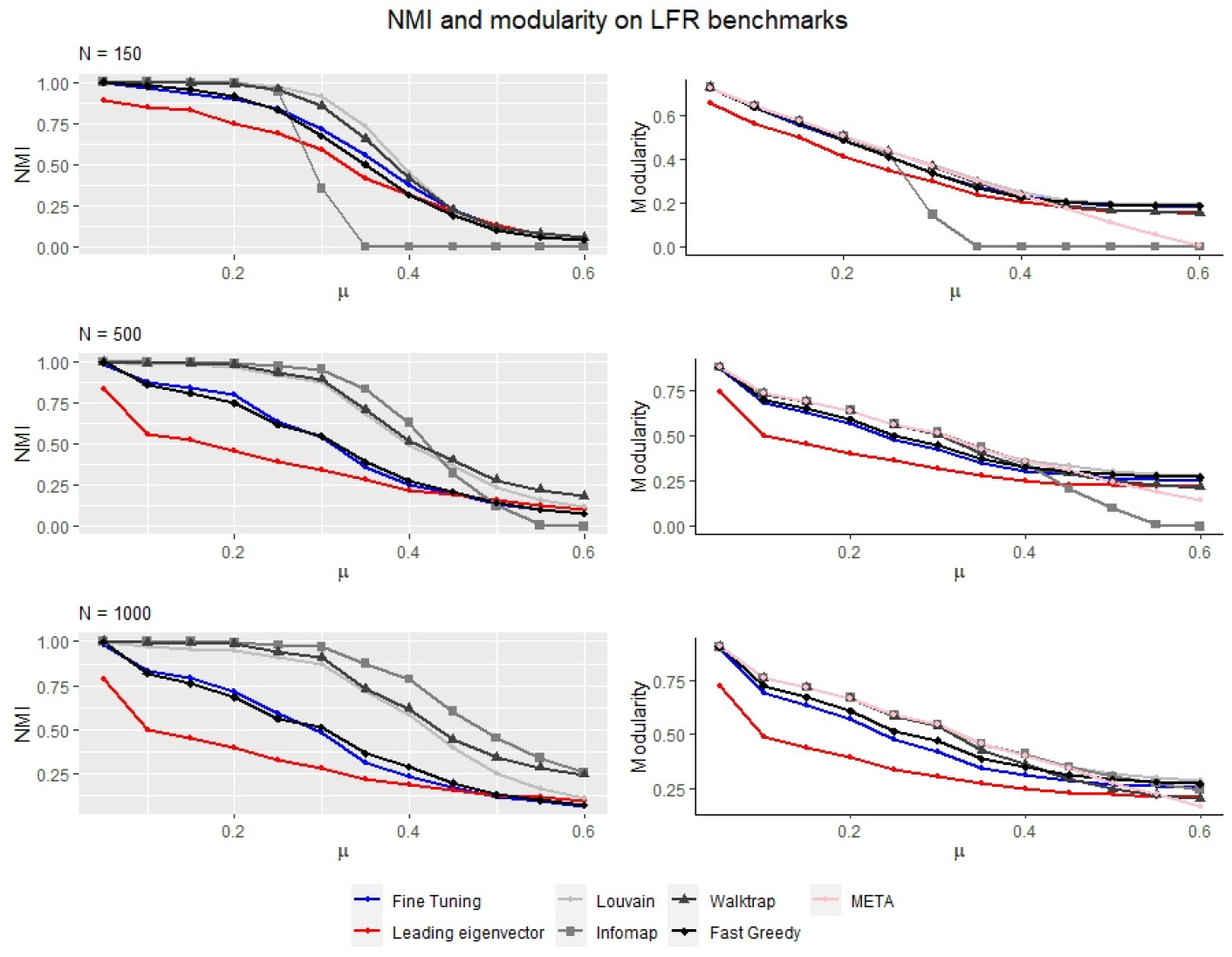
Mean modularity and NMI with planted partitions of the LFR networks.

When we look at the studied waterstrider sample, all the characters are highly correlated: the correlations of antseg4 with other characters are high, while the correlations between the other characters are very high. This pattern emerges because all the characters are largely affected by the overall size. We excluded the effects of sex and instar which resulted in considerably lower correlations, in particular for the antseg4 (Fig. 5), The final correlation matrices were calculated from the transformed data.

**Fig. 5.**
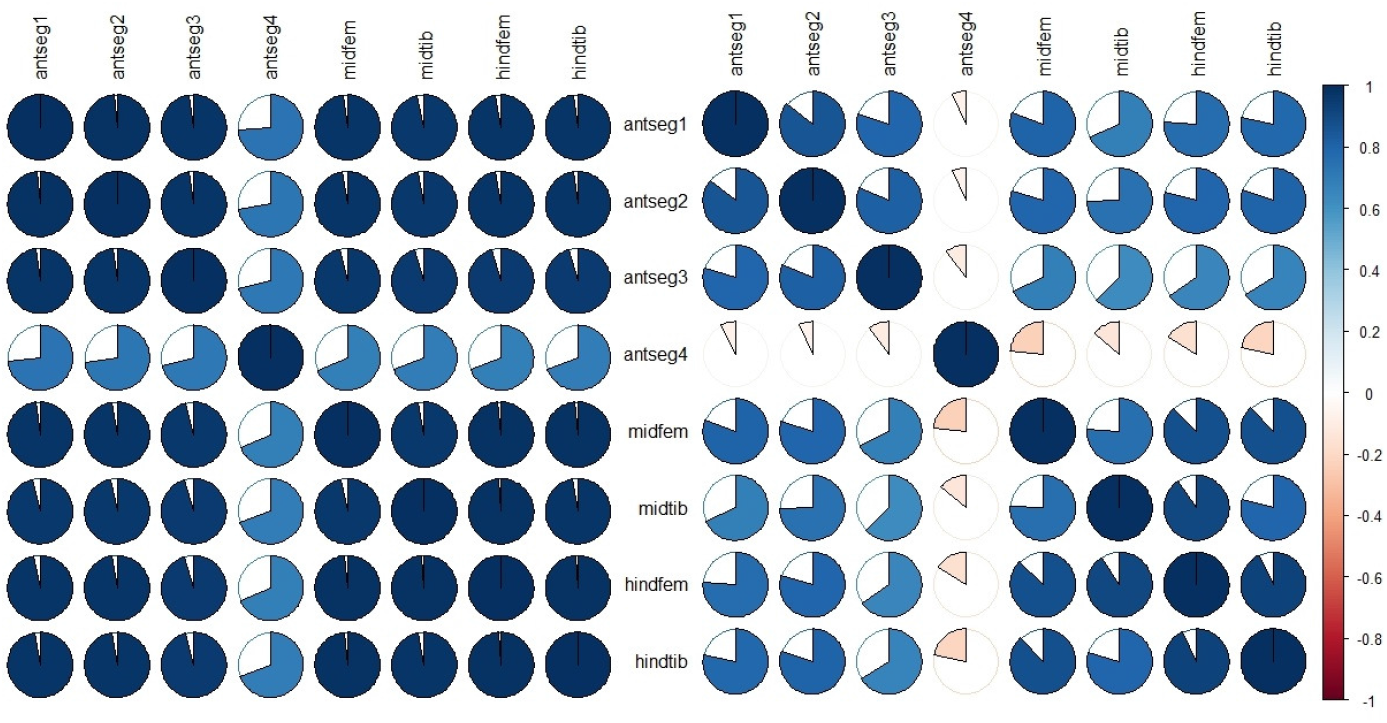
Correlation matrices before and after excluding the effects of sex and instar; antseg 1-4 are the lengths of 1st to 4th antennal segments, others are the lengths of middle and hind femora and tibiae.

**Fig. 6.**
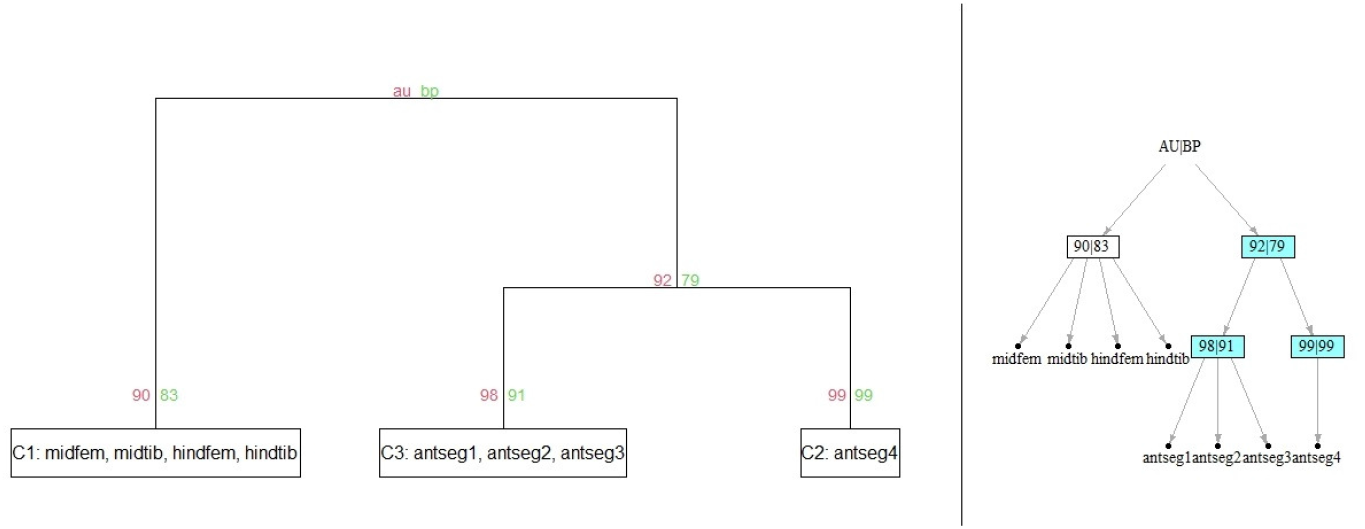
Module structure as depicted by the two functions available: PV.graph (right) and PV.dendro (left). Modules with AU p-values above 90% level are highlighted in the light grey. Abbreviations of characters are in Fig. 5. Correlation coefficients were squared and cut at the 0.15 quantile.

LModularity function with optimal algorithm option gave pattern with all the measurements, except from midtib and hindfem, attributed to separate modules, or correlation pleiades. Such a pattern can be interpreted that all the measurements belong to one module instead.

In comparison, our procedure revealed three modules or correlation pleiades (Fig. 5). The characters of the antennal segments 1-3 and the legs form the 1st and the 3rd pleiades respectively, while one more antennal character, antseg4, is on its own, the 2nd, pleiad (apparently due to lack of correlation with other characters). The result is similar to the result of the *pvclust* function that uses hierarchical clusterization based on the correlation matrix. The AU p-values estimates calculated with *pvclust* also show strong support for the clusters found by the leading eigenvector algorithm.

Such a partition agrees well with the biological information. In Heteroptera, the function of antennae is primarily sensory (e.g., Nowińska & Brożek (2017)), whereas leg segments are the elements of locomotor apparatus. The assignment of the length of the 4th antennal segment to its own pleiad may indicate that this segment is not as closely related to other segments in development or functioning as the other segments are. However, it also may be due to the sampling error; the result deserves checking on a larger sample. Decreasing correlation from proximal to distal elements has been reported for antennae of another heteropteran, *Pyrrhocoris apterus* L., 1758 (Pyrrhocoridae, Heteropteran, Insecta) (Alpatov & Boschko-Stepanenko, 1928), for limbs in some birds (Alpatov & Boschko-Stepanenko, 1928) and mammals (Bader & Hall, 1960). Usually decreasing correlation is combined with one more pattern, namely increasing variation along the proximo-distal axis, sometimes with the exception of tip element (e.g., Bader & Hall (1960); Hallgrímsson et al. (2002); Holliday & Ruff (2001)) but this is not the case of the studied sample of L. notabilis. In some cases the proximo-distal pattern was not detected. Variance and covariance structures tend to be patterned in evolution according to functional and developmental relationships (Berg, 1960; Cheverud, 1982; Hallgrímsson et al., 2002). This patterning is manifested in subdivision of a character set into the correlation pleiades. For mammalian skull, it is shown that although correlation pattern is rather conservative, the levels of morphological integration vary among the taxa (i.g., Porto et al. (2009); Marroig et al. (2009)) and even within the closely related clades (i.g., Dzeverin (2020)). The level of morphological integration seems to be maintained by selection so as to ensure the efficient performance of organs and and organ parts. The effects of correlational selection are quite important for such a process (Svensson et al. (2021)).

## 4 Discussion

There are various methods available to study integration and modularity, which can yield quite different results (Klingenberg, 2014; Goswami & Polly, 2008; Adams et al., 2004). There are also many state-of-art methods and software solutions that use networks and rely on community detection methods. The leading eigenvector together with the fine-tuning procedure is one of the available algorithms to be applied in the research. At the same time, the very problem of modular structure analysis poses great complexity. For example, the “No Free Lunch” theorem shows that no algorithm can be optimal on all kind of inputs (Wolpert & Macready, 1997), and so the methods’ efficacy would differ on different data, and on the other hand, and the so-called “No Ground Truth” theorem states that there is no bijection between network structure and ground truth communities, and therefore no algorithm can always recover the correct ground truth on every network (Lancichinetti et al., 2008). Such limitations point towards the necessity of careful considerations regarding both the methods and final selection. The proposed procedure based on multiscale bootstrap offers an alternative method for discovering modular structure of the data with the ability to assess statistical significance of the result and thus to allow researchers to make an informed decision regarding the final partition. Overall, both methods had lower modularity compared to the other methods on the simulated networks. The difference of the performance of the methods (NMI, modularity) for the LFR benchmark and the real-world *CommunityFitNet* networks might be attributed to the structural differences between the benchmark model and the real-world networks; similar differences have been also suggested in J. Yang & Leskovec (2012) and Orman et al. (2013). The question of the types of networks on which the leading eigenvector algorithm might fail or the ones it is best suited for are of great interest and need further investigation.

## Acknowledgements

We are grateful to Yu. S. Mishura for the idea of preparing an article based on our results. We are grateful to Maria Ghazali and Pavel Gol’din for valuable comments, suggestions, and criticisms. We thank C. P. Klingenberg and J. R. Spence for providing open access to the waterstrider dataset that we used as an illustrative example. We thank P. V. Putchkov for helpful comments. Thanks also are due to the staff of the Department of Probability Theory, Statistics and Actuarial Mathematics of Taras Shevchenko National University of Kyiv for help, advice and support.

## Declarations

### Code availability

The R language code for the procedure described in the article is available in the github repository https://github.com/OVertsim/LVPV.

